# Ecology of hydrocarbon degradation in widespread viruses

**DOI:** 10.64898/2025.12.08.692962

**Authors:** Linyi Ren, Kaiyang Zheng, Yantao Liang, Hongmin Wang, Ziyue Wang, Yundan Liu, Xinran Zhang, Yue Dong, Hongbing Shao, Xiyang Dong, Andrew McMinn, Min Wang

## Abstract

Hydrocarbons are vital for energy production and lead to serious environmental issues due to pollution. Microorganisms largely drive the production and degradation of hydrocarbons, yet little is known about viral contributions to hydrocarbon degradation. Here we identified 786 viral contigs from IMG/VR(v4), encoding five aerobic (*alkB*, *ladA*, *almA*, *ndoB* and *dszC*) and three anaerobic (*bssA*, *ebdA* and *abcA*) hydrocarbon-degrading genes (HDGs). vOTUs encoding HDGs span 26 distinct viral families, including 249 are mainly associated with host-associated environment and 463 are derived from aquatic habitat, respectively, implying that these viruses are broadly engaged in hydrocarbon-degradation processes across the entire biosphere. *alkB* (35%) and *almA* (28%) was tended to be encoded by Schizomimiviridae, *bssA* was tended to be encoded by T5-like bacteriophages (29%) and SPO1-like bacteriophages (22%), suggesting that the carriage of HDGs by viruses exhibits taxonomic specificity. With respect to host associations, Pseudomonadota and Bacillota constitute the two main potential host lineages, associated with 268 and 71 vOTUs, respectively. The profiling of ecological footprint reveals that viruses contribute a total of up to 30% gene abundance and 32% transcribing activity to hydrocarbon degradation in the global ocean. This study systematically revealed the distribution, diversity, virus-host correlation and activity of virus-encoded HDGs, underscoring the significant role of viruses in hydrocarbon metabolism on a global scale.

## Introduction

Hydrocarbons, such as alkane, are vital for energy production and various industrial processes. However, they also lead to environmental issues due to pollution [1]. The combustion of hydrocarbons for energy emits carbon dioxide and other pollutants, contributing to air pollution and climate change [2]. Additionally, accidental spills, leaks, or improper disposal of hydrocarbons can pollute soil and water, harm ecosystems and pose health risks [3, 4].

Microorganisms play a crucial role in the complete degradation of hydrocarbons into CO_2_ and H_2_O [5, 6], offering a permanent solution to eliminate pollutants without causing secondary environmental issues [7–9]. Certain bacteria exhibit the ability to degrade hydrocarbons either aerobically or anaerobically [10–13]. Remarkably, hydrocarbon degradation bacteria, such as *Oleibacter*, *Thalasolitus*, and *Alcanivorax*, thrive even at extreme depths exceeding 10,400 meters within the Mariana Trench [14]. The enzymes involved in the initial steps of hydrocarbon degradation include hydrocarbon hydroxylases (AlkB), flavin-binding monooxygenase (AlmA), long-chain hydrocarbon hydroxylase (LadA), naphthalene-1,2-dioxygenase (NdoB), benzylsuccinate synthase (BssA), dibenzothiophene monooxygenase (DszC), molybdopterin-family ethylbenzene dehydrogenase (EbdA), and benzene carboxylase (AbcA) [15]. The genes encoding these hydrocarbon-degrading enzymes demonstrate diverse capabilities, allowing them to efficiently break down various types of alkanes or aromatic hydrocarbons under different environmental conditions [16]. The diversity of these genes and the vitality of hydrocarbon-degrading microorganisms highlight the potential for natural restoration of hydrocarbon pollution across various ecosystems.

Viruses, the most abundant and diverse “biological entities” on the Earth, exert influence on host communities by infecting and lysing host cells. They manipulate host metabolic pathways through the expression of virus-encoded putative auxiliary metabolic genes (AMGs), indirectly impacting global biogeochemical cycles [17].

Additionally, viruses play a crucial role in horizontal gene transfer (HGT) within the microbiome, mediating gene recombination [18–20]. Advances in metagenomic techniques have opened avenues for exploring the diversity and ecological functions of viruses encoding essential AMGs in various environments [21]. Despite the importance of hydrocarbon degradation in biogeochemical cycles, the viral communities that interact with these microbial players, especially in the context of hydrocarbon degradation, remain poorly characterized. Studies have shown that large-genome phages infecting methanotrophs carry the *pmoC*, and may influence methane oxidation and carbon cycling[22]. And the study by Ru et al. primarily focuses on the diversity and biogeographical distribution of virus encoded six alkane degradation genes, and highlights how phage AMGs contribute to the bioremediation of hydrocarbon pollution and promote the proliferation of these microorganisms[23]. Although hydrocarbon degradation AMGs have been identified in multiple hosts or viruses, their interactions and roles in the biogeochemical cycling of hydrocarbons remain poorly understood. Characterizing the ecology, function, and virus-host interactions of hydrocarbons degradation-related viruses is crucial for a comprehensive understanding of the mechanisms of hydrocarbon degradation and metabolism.

Here, through analysis of publicly available metagenomic databases, we identified 786 viruses encoding hydrocarbon degradation genes (HDGs) across aquatic, host-associated and engineered environments. Our assessment reveals the distribution, diversity, potential host, and putative activity of these viruses, emphasizing the essential need to acknowledge and incorporate the contributions of viruses in hydrocarbon degradation. This study emphasizes the significance of viruses as active participants in hydrocarbon metabolism across diverse ecosystems, reveal their ecological roles and provide a comprehensive understanding of hydrocarbon degradation processes.

## Results

### Distinct viruses encoding AMGs for the degradation of hydrocarbons and aromatic hydrocarbons

Based on the HMM models from the CANT-HYD, we searched IMG/VR(v4), which includes over 15 million virus genomes and genome fragments, and identified 768 viruses encoding hydrocarbon-degrading genes (VHDGs) (Figure 1A and Supplementary Table 1)[24, 25]. Among them, 554 are classified as genomic fragments, 145 as high-quality viruses, and 69 as reference sequences. The viruses encoding *alkB* were the most prevalent (n=275), primarily involved in the degradation of C5-C13 alkanes. Following *alkB*, *bssA* is the second most abundant hydrocarbon-degrading genes (HDGs) encoded by viruses (n=233), engaged in the anaerobic degradation of aromatic hydrocarbons (Figure 1A). The viruses encoding *ladA* (*ladA-*viruses, 89.4%), *alkB* (*alkB-*viruses,98.9%), *ndoB* (*ndoB-*viruses,100%), *dszC* (*dszC-*viruses, 84.8%), and *almA* (*almA-*viruses, 96%) are predominantly distributed in aquatic, where aerobic hydrocarbon degradation is more prevalent. These genes are involved in the degradation of various hydrocarbons through aerobic pathways. In contrast, viruses encoding *bssA* (*bssA-*viruses, 61.8%) and *ebdA* (*ebdA-*viruses, 53.5%) and *abcA* (*abcA-*viruses, 50%), which are involved in anaerobic hydrocarbon degradation, are mainly found in host-associated environments (Figure 1B).

**Figure 1.**
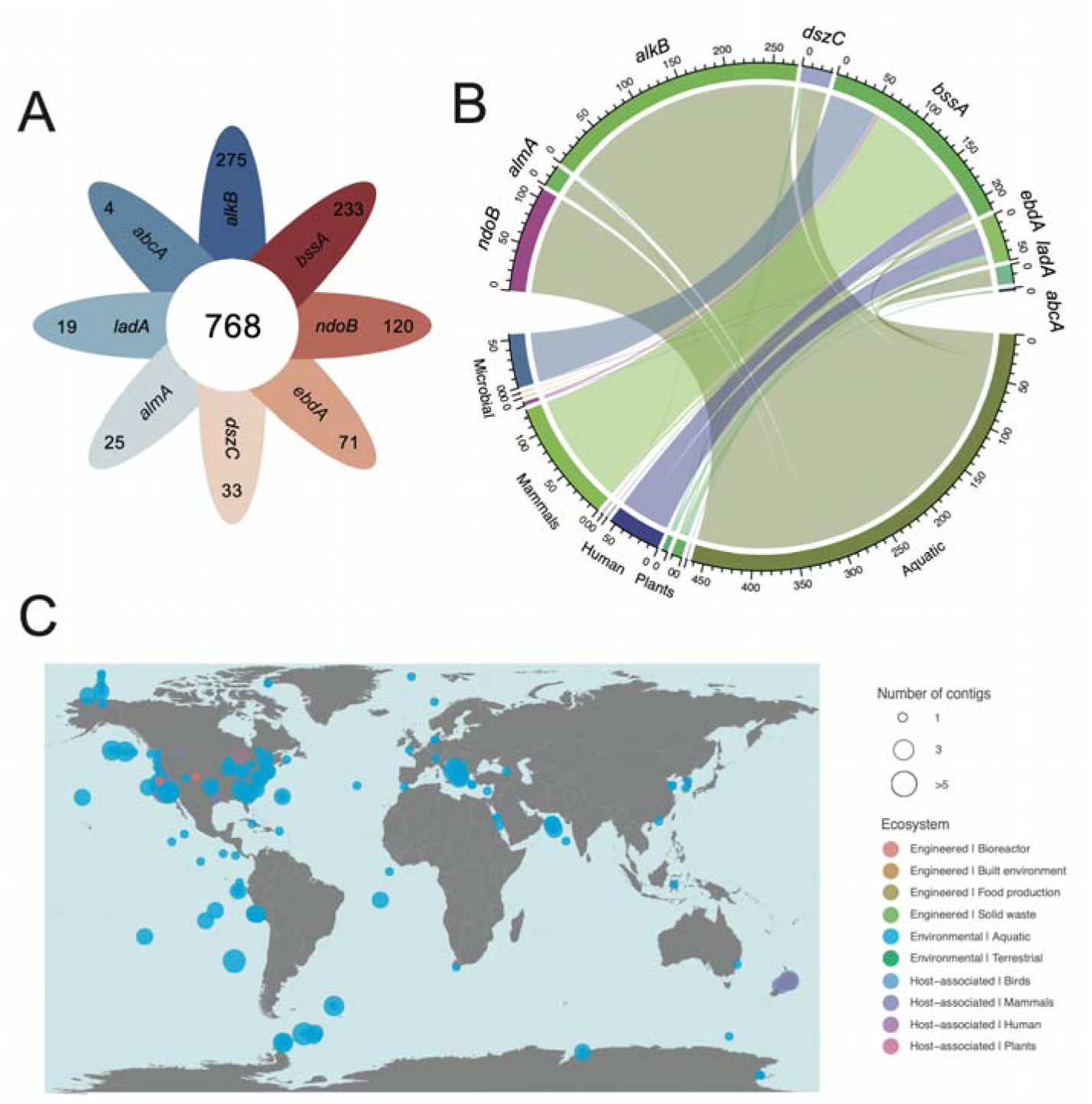
Distribution of metagenomic viruses encoding HDGs in global ecosystems. (A) Venn diagram illustrating the distribution of VHDGs across different categories. The central circle represents the total number of VHDGs (768), with each surrounding segment corresponding to a specific VHDG category. The numbers within each segment indicate the quantity of each VHDG: abcA (4), ladA (19), almA (25), dszC (33), alkB (275), bssA (233), ndoB (120), and ebdA (71). (B) Circos plot illustrating the distribution of VHDGs across different environments. Each outer segment represents a specific VHDG and environment, and the connections within the plot indicate the habitats in which the corresponding viral sequences were found. (C) The distribution of VHDG in global scale. Each VHDG-detection metagenome is represented by a circle proportional to the number. The different habitats were labeled by different colors.

The VHDGs generated in this study came from 440 different metagenomic

samples, most of which were from aquatic (n=351), followed by host-associated (n=74) and engineering (n=14) (Figure 1C). HDGs encoded by viruses varied across ecosystems, with aquatic predominantly featuring *alkB*, *ndoB*, *dszC*, *almA* and *ladA* involved in aerobic hydrocarbon degradation. Conversely, in host-associated ecosystem, the HDGs encoded by viruses are *bssA*, *ebdA*, and *abcA*, which are involved in the anaerobic degradation of hydrocarbons. This result indicates the hydrocarbon-degrading genes have an impact on the ecosystem of viruses (Figure 1C).

### Initial degradation mechanism of hydrocarbons by VHDGs

In this study, molecular docking was employed to analyze the individual interactions between VHDGs and the ligand hydrocarbons, in order to determine the affinity of these HDGs for hydrocarbons at their catalytic sites. The binding free energy is stable when the value is negative, which can be understood as under isothermal and isobaric conditions, after the receptor and ligand are bound, the system becomes more stable[26]. In this study, representative VHDGs were selected and subjected to molecular docking with the substrate. Figure 2 presents the docking structure of the hydrocarbon-enzyme complex, illustrating the docking positions and highlighting the amino acid residues surrounding the optimal binding site for the alkane (Figure 2). For BssA and EbdA, which are involved in aromatic hydrocarbon degradation under anaerobic conditions, the binding energies with the substrate are -4.24 Kcal/mol and -4.75 Kcal/mol, respectively. These binding energies are similar to those of the bacterial HDGs in relation to substrate[26]. Additionally, toluene exhibits hydrophobic interactions with VAL, ASN, TYR, PHE, and ALA of BssA, which is consistent with previous studies on the active site of BssA[27]. Ethylbenzene shows hydrophobic interactions with ILE, PRO, and PHE of EbdA. Dibenzothiophene (DBT) binds between PHE, ILE, and GLN of the viral-encoded DszC, with a binding energy of -6.04 Kcal/mol. It is known that PHE is located at the entrance of the substrate binding site and directly interacts with DBT, while GLN and ILE are also present in the active site of DszC[28, 29]. Similarly, NdoB is also a gene that degrades aromatic hydrocarbons under aerobic conditions, and its binding energy is -4.29 Kcal/mol. For HDGs involved in alkane degradation, octane interacts with hydrophobic amino acids such as ALA, TYR, and ILE in the viral-encoded AlkB, with a binding energy of -4.25 kcal/mol. Studies have shown that the hydrophobic input channel is the preferred region for AlkB to bind hydrocarbons, and the polarity and negative charge of GLU may play a role in helping to position or stabilize the substrate[30, 31]. The 2-hexadecanone and the Ile, Leu, Trp, Phe, and Val in virus-encoded AlmA form a hydrophobic environment, possibly to accommodate the hydrophobic aliphatic chains of LC ketones[32]. The presence of Tyr and Lys may play a role in stabilizing the structure or modulating the interaction. Similarly, the binding energy of pentadecane to viral LadA is -4.32 Kcal/mol, similar to previous studies on Aspergillus flavus[33]. These findings suggest that the viral-encoded hydrocarbon-degrading genes (HDGs) exhibit significant binding affinities for various hydrocarbons, indicating their potential role in hydrocarbon degradation. This reinforces the hypothesis that viral HDGs possess the enzymatic capability to degrade hydrocarbons, similar to bacterial counterparts.

**Figure 2.**
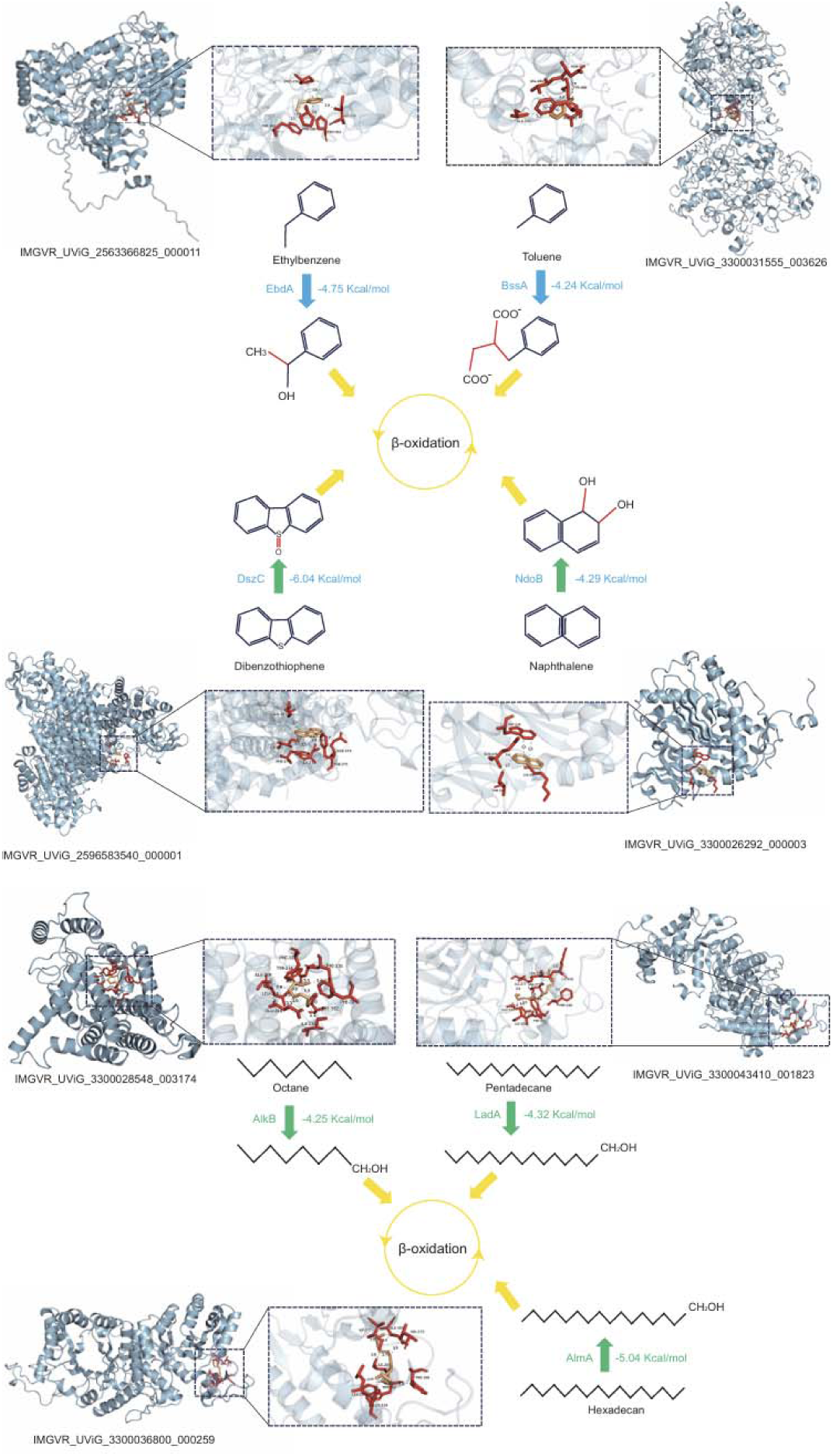
Key reaction steps, molecular docking results and binding energys for the degradation of aromatic compounds. Gene names: AlkB, hydrocarbon hydroxylases; AlmA, flavin-binding monooxygenase; LadA, long-chain hydrocarbon hydroxylase; NdoB, naphthalene-1,2-dioxygenase; BssA, benzylsuccinate synthase; DszC, dibenzothiophene monooxygenase; EbdA, molybdopterin-family ethylbenzene dehydrogenase.

### VHDGs are taxonomic diverse

We applied VITAP to taxonomically classify and cluster the identified VHDGs[34]. The majority of VHDGs were assigned to Caudoviricetes (n=540, ∼70%) and Megaviricetes (n=214, ∼28%). At the order level, the most annotated Megaviricetes VHDGs is *Imitervirales* (n=194, ∼91%), and most of the VHDGs in Caudoviricetes is unknown (Figure 3A).

**Figure 3.**
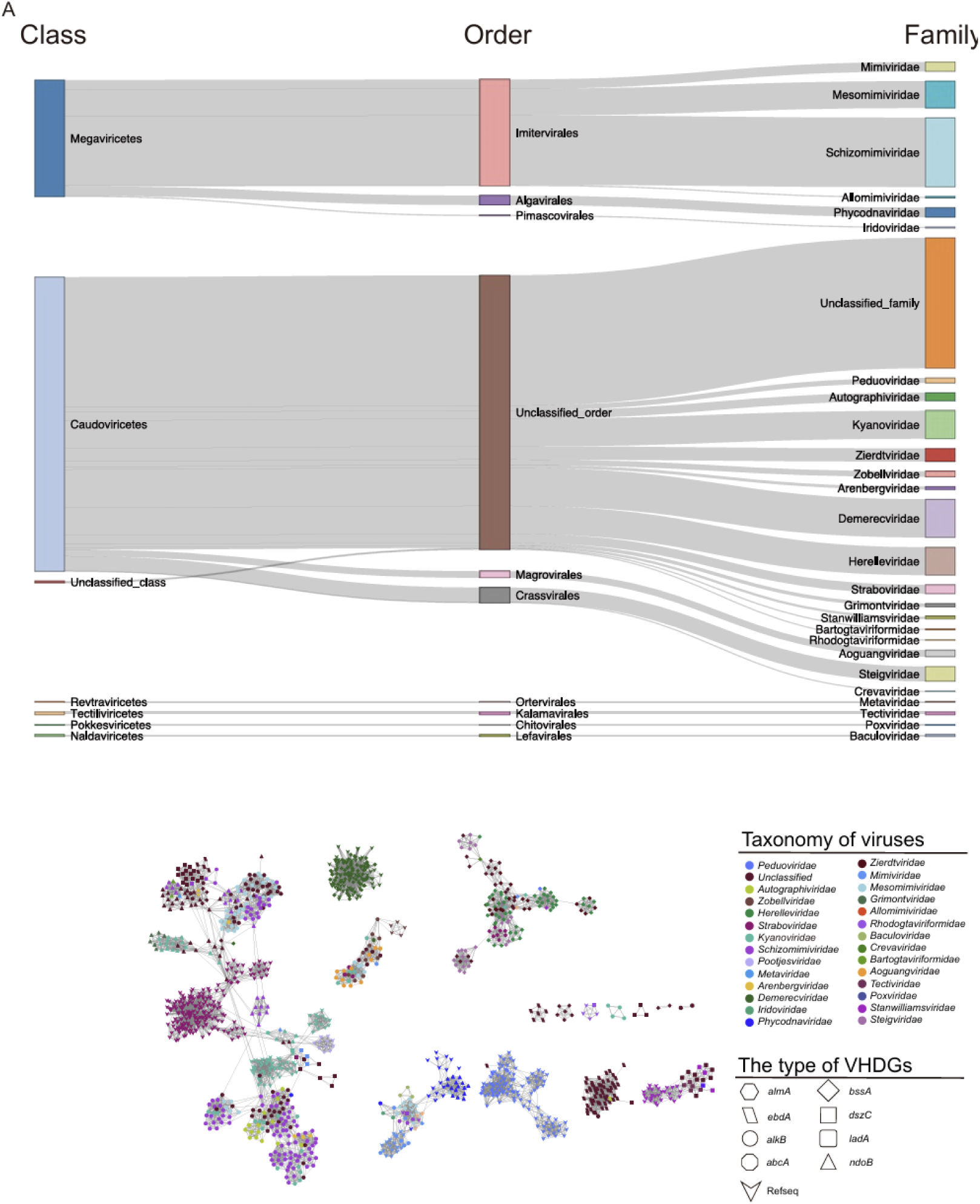
Taxonomic assignment of metagenomic VHDGs and protein network clustering with reference viruses. (A) The proportion of VHDGs at the class, order and family level. (B) Gene-sharing network of VHDGs and RefSeq from ICTV. Viruses (nodes) are connected by edges, indicating the significant pairwise similarity between them in terms of shared protein contents. The different colors of the dots represent different viral taxonomy, and the shapes represent the types of HDGs encoded by the virus.

In accordance with these results, we constructed a protein sharing network of the VHDGs with reference viruses from the NCBI GenBank database (Figure 3B). *bssA*-viruses are associated with the widest range of viral taxa, with viruses from 13 different families found to encode *bssA*. They are mainly divided into two groups. One group is entirely composed of *bssA*-viruses, primarily belonging to Herelleviridae, Steigviridae, and Zierdtviridae. The other group consists of *bssA*-viruses reference viruses from Demerecviridae. *alkB* -viruses are associated with 12 different viral families and primarily divided into three groups in the network. They mainly belong to *Schizomimiviridae*, *Autographiviridae*, and *Mesomimiviridae*. *ebdA*-viruses cluster into a single group and remain unclassified at the family level. *ebdA*-viruses are associated with the fewest range of viral taxa, only a small number classified into two known viral families, while the majority belong to unclassified viruses. Viruses encoding *ndoB* are primarily divided into three groups. In the first group, *ndoB*-viruses cluster with reference sequences belong to *Phycodnaviridae*. In the second group, these viruses cluster with *dszC*-viruses and belong to *Schizomimiviridae* and unclassified viruses at the family level. In the third group, *ndoB*-viruses cluster separately and belong to *Kyanoviridae* and *Grimontviridae*. Viruses encoding *lada* cluster with reference sequences and belong to *Tectiviridae* and *Stanwilliamsviridae*. Overall, viruses encoding the same HDGs tend to cluster together and are associated with a broad range of viral taxa, which may suggest the widespread presence of viruses carrying HDGs in the environment.

### Virus-host linkages and viral lifestyles

To explore virus-host interaction of VHDGs, potential hosts were predicted for 768 VHDGs using iPHoP[35]. As a result, 1743 virus-host linkages were predicted, more than half of the VHDGs predicted the potential host (n=402, ∼53%). Most of these VHDGs infect Pseudomonadota (n=249, ∼65%), which is widely present in various environments and facilitating both aerobic and anaerobic hydrocarbon degradation. The primary potential host for the viruses encoding *bssA* is Bacillota (n=56, ∼64%), which differs from other hydrocarbon-degrading gene viruses (Figure 4A). Research has shown that this is also the primary bacterial lineage encoding *bssA* [26]. A total of 12 bacterial phyla were predicted as hosts for VHDGs, underscoring the broad host range and ecological diversity of these viruses. Through a review of the literature, we found that VHDGs infect known hydrocarbon-degrading bacteria (Figure S1). At the genus level, the most prevalent result was *Escherichia* (n=63, ∼16%), which is a bacterial group capable of degrading both aliphatic and aromatic hydrocarbons[36, 37]. The primary viruses infecting these bacteria are *ebdA*-viruses (n=56, ∼86%). They also infect *Enterobacter* and *Klebsiella* of *Enterobacteriaceae*, as well as *Xanthobacteraceae* bacteria, all of which have been demonstrated to possess aromatic hydrocarbon-degrading abilities[38–40]. In addition, we found that viruses encoding *bssA* also infect *Escherichia* and *Enterobacter*. However, viruses encoding other HDGs exhibit a distinctly different host range, including *Pseudoalteromonas*, *Acinetobacter*, *Vibrio*, *Flavobacteriaceae*, *Burkholderiaceae*, and others[41–45]. The involvement of multiple bacterial phyla, each with distinct metabolic capabilities and environmental niches, suggests a complex and widespread network of viral-host interactions facilitating hydrocarbon degradation across diverse habitats.

**Figure 4.**
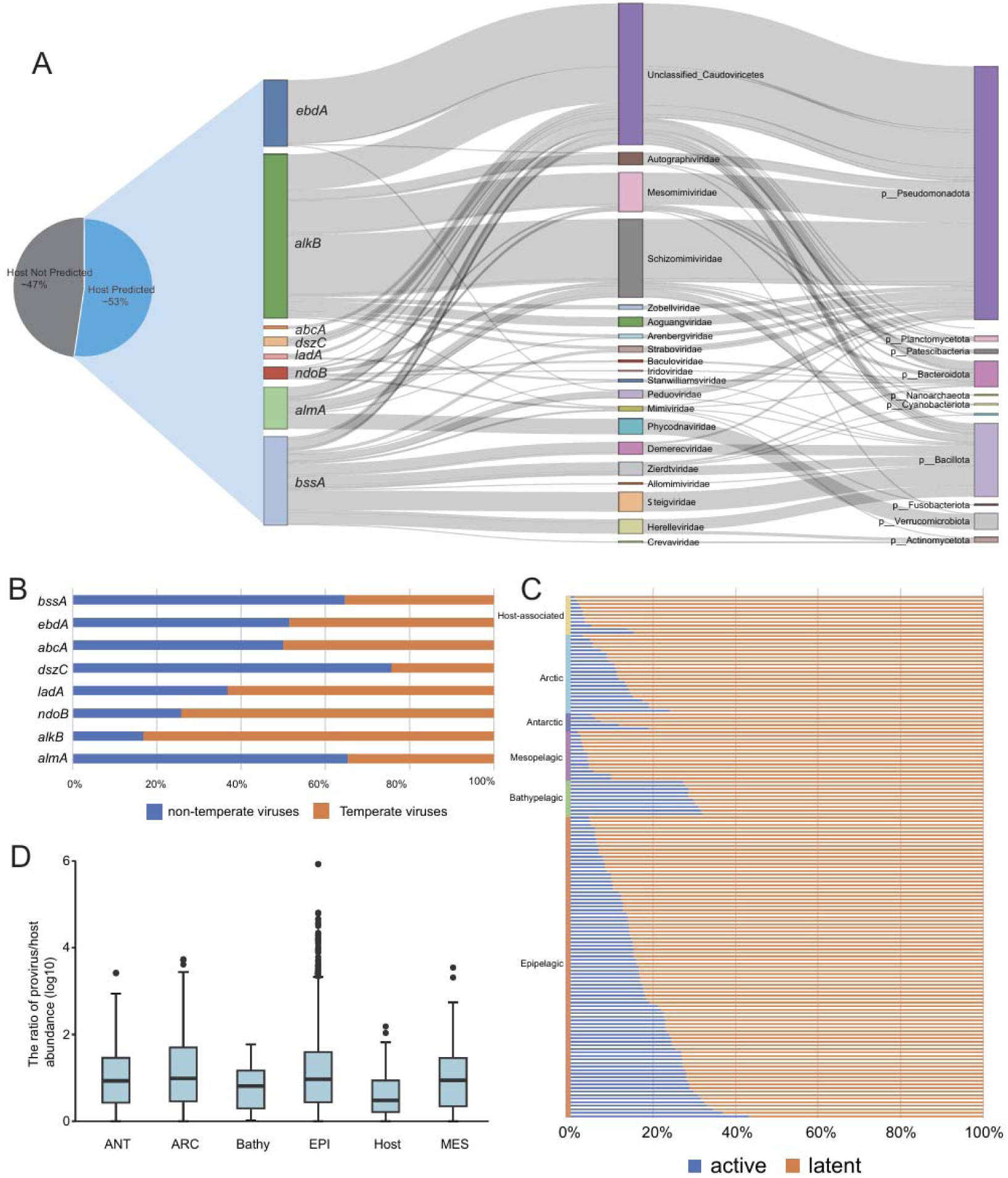
Predicted host-virus interactions. (A) Predicted virus-host linkages. Percentage and taxonomy of VHDGs for which a host was predicted are shown on the left; the taxonomy of predicted hosts is shown on the right. (B)Lifestyle of viruses encoding different types HDGs. (C) Active lysogenic VHDGs in ocean and host-associated environment. The percentage represents the proportion of potentially active VHDGs in lysogenic VHDGs. (D) Provirus/host abundance ratios of ocean zone (epi-/mes-/bathy/ant/arc) and host-associated. The ratios were calculated by the average per-base-pair coverage of the provirus and host contigs, respectively. EPI: Epipelagic; MES: Mesopelagic Zone; Bathy: Bathypelagic; ANT: Antarctica; ARC: Arctic.

To explore the lifestyles of VHDGs, we identified 349 proviruses. The number of prophages among viruses encoding different types of HDGs varies significantly. Notably, prophages were most abundant in viruses encoding the *dszC* (n=25, ∼76%), *alma* (n=17, ∼65%) and *bssA* (n=154, ∼65%), while those encoding the *ndoB* (n=31, ∼25%) and *alkB* (n=47, ∼16%) found fewer prophages (Figure 4B). Since most of the VHDGs in this study are derived from aquatic (n=463,∼60.3%), we calculated the provirus-host abundance ratio (PVHR) in GOV2.0[46, 47]. Results showed that a considerable proportion of proviruses in Bathypelagic (average=30%) were potentially active, while the proportion of potentially active proviruses discovered in the Mesopelagic (average=4%) and host-associated environment (average=5%) is the lowest. In addition, the proportion of potentially active proviruses varies significantly among different sampling stations in Epipelagic (range=38%) and Arctic (range=21%) (Figure 4C). Further analysis found that PVHR was similar across aquatic and higher than in host-associated environment, suggesting a high production rate or burst size of proviruses in aquatic (Figure 4D). In particular, IMGVR_UViG_2563366825_000011 that was predicted to integrate into the genome of GCF_000008865.2, which showed a high abundance in Bathypelagic, Arctic and Epipelagic (Figure S2). GCF_000008865.2 belongs to Escherichia, which is also a hydrocarbon-degrading microorganism. Through HMM search, it was found that it encodes *dszC*, *ebdA*, *ndoB*, *nmsA*, and *ladA*. Therefore, active IMGVR_UViG_2563366825_000011 may play an important role in regulating microbial hydrocarbon degradation. Collectively, lysogenic infection appears to be the preferred lifestyle for VHDGs in marine environments, particularly in the deep sea.

### The evolutionary uniqueness of viral HDGs

Phylogenetic inference by maximum likelihood is widely used in molecular systematics[48]. Here we carried out a comprehensive search for HDGs from bacteria and archaea derived from the reference sequences of NCBI (https://ftp.ncbi.nlm.nih.gov/genomes/refseq/). We identified HDGs in these contigs using a broad-spectrum hidden Markov model (HMM) profile and subsequently selected non-redundant HDGs protein sequences (similarity <90%; Supplementary Table 1). The AlkB, NdoB, BssA, EbdA can be divided into three groups on phylogenetic tree, respectively. The vast majority of the virus-encoded AlkB is in group3, showing obvious monophyletic characteristics, suggesting that these viruses share a common evolutionary ancestor and exhibit high genetic similarity within this group. Similar to AlkB is virus-encoded EbdA, which is mostly concentrated in group2 and closely related to bacterial-encoded HDGs. The virus-encoded NdoB and BssA are primarily distributed in groups 2 and 3 of their phylogenetic trees, and both exhibit strong monophyletic characteristics (Figure 5). The number of AlmA, LadA, and EbdA encoded by viruses is relatively small, and most of these genes co-evolve with HDGs encoded by bacteria (Figure S3). In addition, the dendrogram based on the HDGs structure exhibits a topology similar to that of the HDG phylogenetic tree. In particular, viral-encoded AlkB, BssA, NdoB, and EbdA show high structural similarity and cluster independently (Figure S4). The phylogenetic signal of these proteins not only mirrors the substantial diversity of HDGs but also uncovers previously uncharacterized monophyletic HDGs encoded by viruses (especially *alkB*, *ndoB*, *bssA*, and *ebdA*). We hypothesize that HDGs encoded by viruses may have originated from bacteria or archaea and evolved distinct characteristics over time.

**Figure 5.**
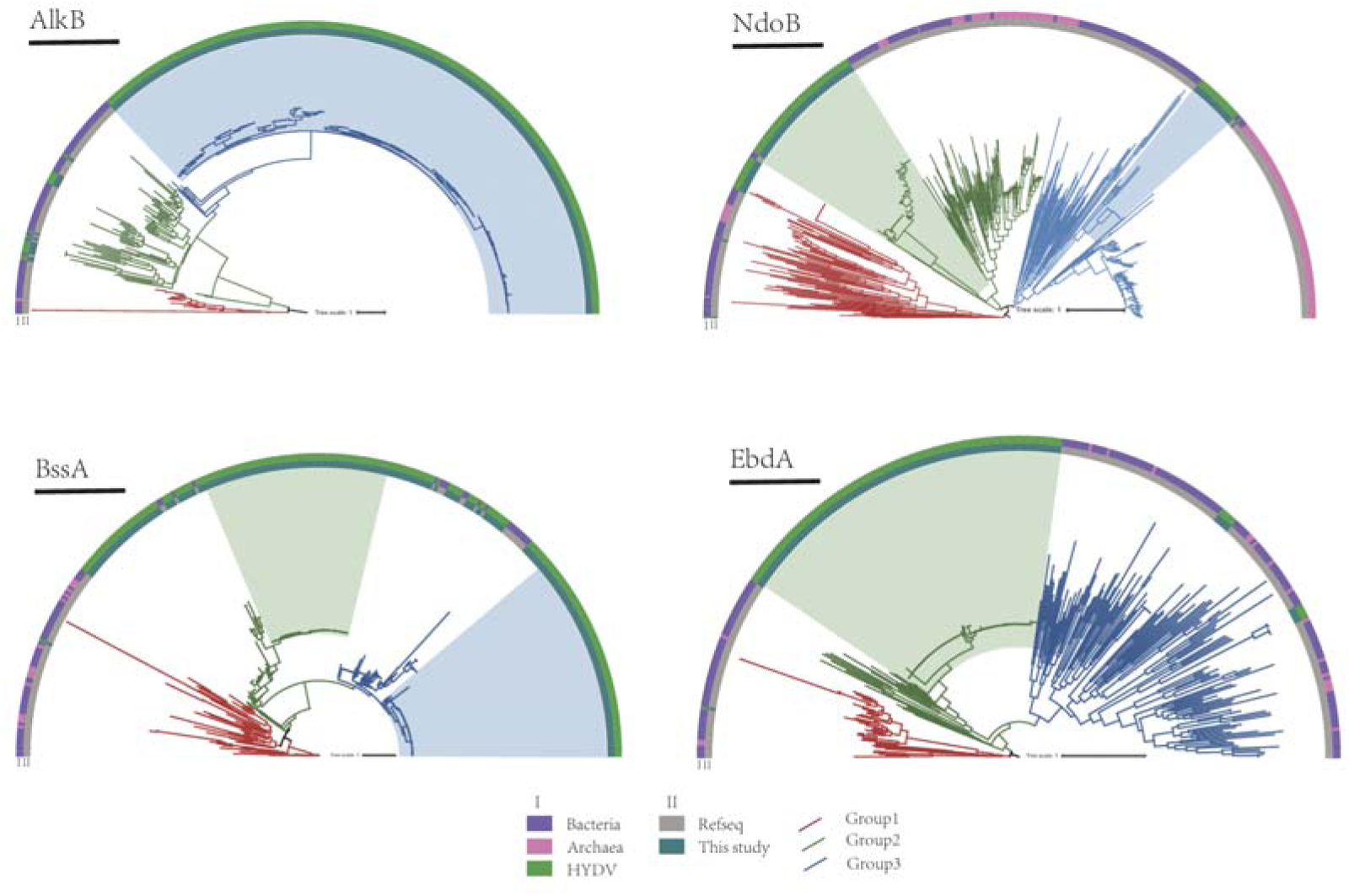
Phylogenetic trees between Bacteria, Archaea and Virus encoded HDGs (Hydrocarbon degradation genes). Posterior mean sites frequency mixed model (LG+C30+F+R10) was applied to each hydrocarbon degradation gene, with the tree rooted at the midpoint. The outer ring represents the origin of HDGs, while the inner ring distinguishes between sequences generated in this study and reference sequences.

### Viral contributions to hydrocarbon degradation based on meta-omic analyses

Based on previous research, we utilized metagenomic datasets to calculate the ratio of virus:total genes for each HDG[49]. The virus:total gene ratios within a community and for each predicted phage-host pair can be used to estimate virus contributions to hydrocarbon degradation. This relies on the assumption that gene ratios can proportionally reflect real metabolic activities, and that host cells in a virocell state retain the same level of environmental fitness as compared to uninfected microorganisms[49].

By mapping metagenomic reads to HDGs encoded by viruses and putative bacterial hosts, we obtained the viral HDGs to total gene ratios, which represents the relative contribution of HDG functions to the representative metabolism. Since most VHDGs are derived from aquatic, we identified virus-host HDGs pairs based on the host prediction and calculated HDGs coverage ratios in Tara Ocean metagenomic datasets and Mariana Trench metagenomic datasets (Figure 6A). Our results show that the average virus:total gene coverage ratios differ in HDGs. At different depths, the *alkB*:total ratio is similar (SRF=20%, DCM=25%, MES=17%), and relative contribution of virus-encoding *alkB* has been found at 10400 in the Mariana Trench (30%). The *ndoB*:total ratio is the highest among all HDGs (SRF=25%, DCM=26%, MES=25%), the relative contribution is stable at various depths. But no contribution of the virus-encoding *ndoB* has been detected in the abyss. The *bssA*:total ratio remains comparable between the surface layer (16%) and the deep chlorophyll maximum layer (18%), but it significantly decreases in the mesopelagic zone (8%). Compared with other HDGs, the *dszC*:total ratio (SRF=9%, DCM=10%, MES=6%) and the *ebdA*:total (SRF=3%, DCM=3%, MES=2%) ratio are relatively low in the ocean. In conclusion, although gene abundance ratios do not necessarily represent function contributions, this scenario still provides a reasonable estimation to suggest considerable hydrocarbon degradation contributions of virus HDGs in virocells.

**Figure 6.**
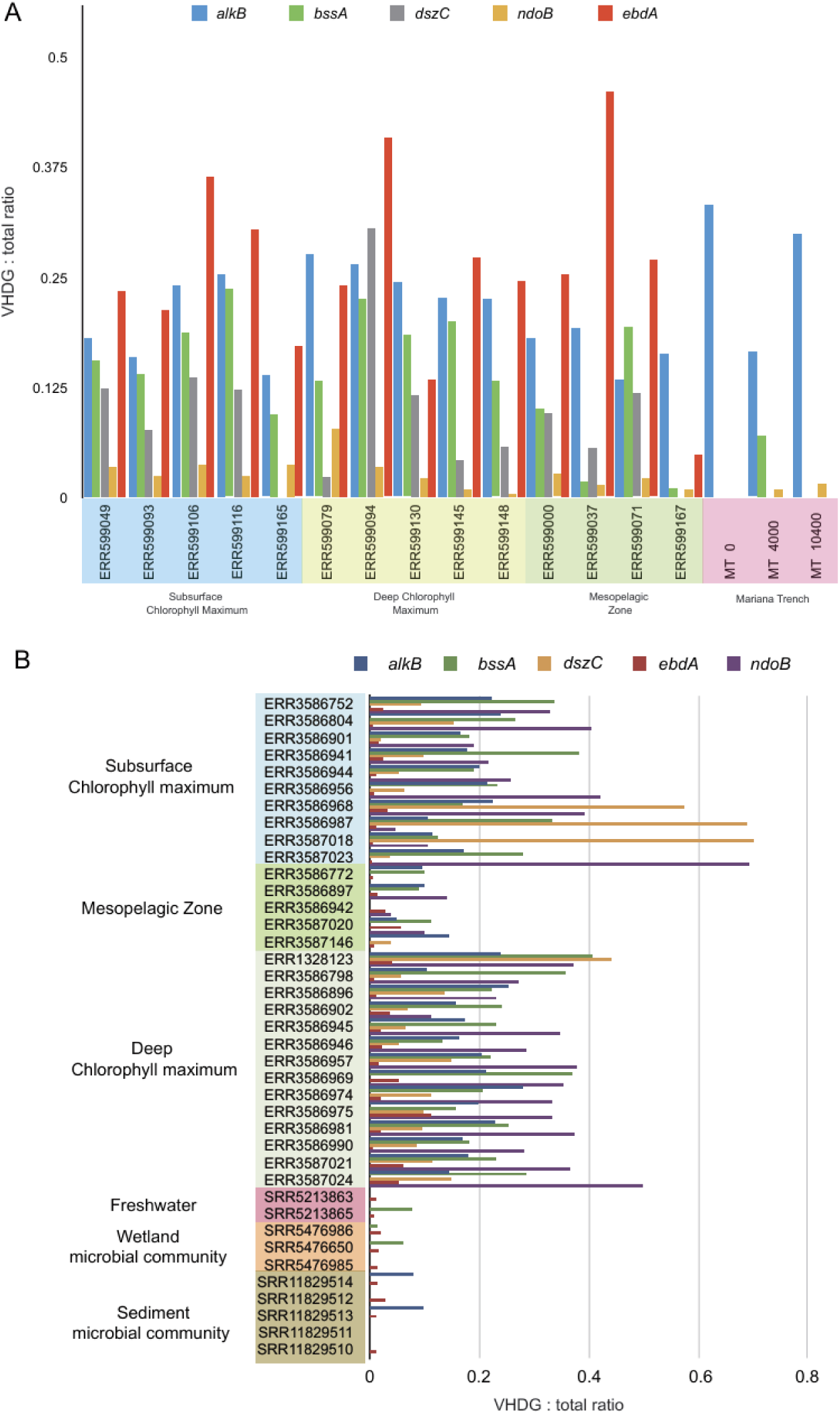
Potential contribution of viral HDGs to their hosts. (A) Viruses HDGs to total (phage and bacterial HDG together) coverage ratios in different depths of the ocean. Different types of HDGs are represented by different colors. (B) Comparison of HDGs expression between virus and host pairs across different ocean depths. Different types of HDGs are represented by different colors.

Subsequently, we used transcriptomic profiling to study HDGs expression in virus:host pairs recovered from Tara Ocean, methane-enriched sediment, wetland, freshwater. The expression ratio of HDGs, measured in RPKM (Reads Per Kilobase per Million mapped reads), exhibit significant variations across different ecosystem and marine depths (Figure 6B). Notably, *ndoB* reaching the highest virus:total ratio (30%) at SRF. The virus:total ratio of *bssA* and *dszC* is similar (25 %), while *alkb* is relatively slightly lower (18%). In DCM, the expression virus:total ratio of *ndoB* (32%), *bssA* (25%) and *aklB*(19%) is similar to SRF, while *dsz*C shows a significant decrease (12%). In MES, all virus:total ratios have shown an apparent decline, especially *dszC*, which has a ratio of only 0.7%. Conversely, the virus:total ratio of *ebdA* exhibits low across all depths, averaging at 2%. In freshwater and sediment, the virus:total ratio of HDGs is very low (<10%), indicating that while these genes are present, their activity is much lower than in the ocean. The above analysis indicates that virus-encoded HDGs may make a significant contribution to host-driven hydrocarbon degradation in the ocean, and the contribution ratios of HDGs vary greatly in different regions and depths.

## Discussion

### Prevalence of rate-limiting enzymes in viral AMGs

The role of viral AMGs in enhancing the degradation of hydrocarbons has been largely attributed to their ability to target and accelerate rate-limiting enzymatic steps within microbial metabolic pathways. These genes, often involved in critical biochemical processes such as the hydroxylation reaction converts methane to methanol, play a pivotal role in microbial communities by alleviating bottlenecks in energy and carbon cycling[50, 51]. Our findings suggest that viruses may facilitate the efficient breakdown of these compounds by encoding enzymes such as *alkB*, *bssA*, and *ndoB*, which are essential in catalyzing steps that are typically slow or rate-limiting(Figure 2). This gene-directed metabolic acceleration is a key feature of viral-host interactions, with significant implications for biogeochemical cycles in hydrocarbon-rich environments.

### Widespread distribution of HDGs across virus taxa

Metagenomic surveys of viral genomes reveal that HDGs are broadly distributed across diverse viral taxa (Figure 1B). The biogeographical distribution and species diversity distribution of these genes, coupled with their presence in rapidly evolving virus genomes, suggests a widespread conservation of these genes within viruses, likely driven by the adaptive advantages conferred to the viruses by encoding such genes[52, 53]. These HDGs may enable viruses to replicate more efficiently and enhance their adaptability to fluctuating ecological niches, supporting the hypothesis that the persistence of these genes across viral lineages provides significant evolutionary benefits. This broad distribution further underscores the essential role of viral-mediated processes in influencing ecosystem dynamics, particularly in environments impacted by hydrocarbon contamination.

### Extensive host range and lifestyle of VHDGs

The versatility of virus-host interactions is exemplified by the broad host range of VHDGs, which can infect a variety of bacteria across different taxonomic groups, especially within the *Pseudomonadota*. This bacterial group is notable for its diverse metabolic capabilities, including the aerobic and anaerobic degradation of hydrocarbons[54]. Our study identified 768 VHDGs, which were associated with 1743 virus-host linkages, highlighting the widespread and diverse nature of these interactions (Figure 4A). The ability of VHDGs to infect multiple bacterial species and mediate horizontal gene transfer may further contributes to the metabolic plasticity of microbial communities, enhancing their ability to degrade complex hydrocarbons efficiently. This characteristic is particularly crucial in environments where rapid adaptation to changing conditions is essential for microbial survival and ecological function.

Lysogenic infection is a prevalent strategy among VHDGs with a significant proportion of prophages identified, particularly in viruses encoding genes such as *dszC*, *almA*, and *bssA* (Figure 4B). This suggests that lysogeny is a common mechanism, enabling VHDGs to integrate into the host genome and ensuring their long-term stability within host populations. This strategy contributes to the persistence of HDGs in microbial communities, particularly in hydrocarbon-impacted environments, enhancing community resilience[55].

Provirus activity is notably influenced by environmental factors, with a higher proportion of active proviruses observed in bathypelagic zones compared to mesopelagic and host-associated environments (Figure 4D). Deep-sea conditions, characterized by stable environmental factors and the availability of hydrocarbon substrates, provide an optimal niche for VHDGs activity[56]. In contrast, provirus activity in epipelagic environments is more variable, influenced by factors such as nutrient availability, temperature, and microbial community composition[57]. These findings highlight the role of environmental context in shaping viral-host interactions, as previously demonstrated in similar ecological studies

### Ecological impact of VHDGs on hydrocarbon degradation

VHDGs demonstrate significant ecological impact through their role in hydrocarbon biogeochemical cycling, as evidenced by their abundance in metagenomic and metatranscriptomic datasets. The analysis revealed that viruses encoding key hydrocarbon degradation genes, such as *alkB*, *bssA*, and *ndoB*, contribute to 18%-30% of associated degradation pathways, underscoring the often overlooked role of viruses in modulating hydrocarbon degradation (Figure 6). While the majority of viral-host interactions occur independently of AMGs, these viruses actively influence microbial metabolism related to hydrocarbon degradation[58]. By calculating the coverage and expression of HDGs in virus-host pairs, the results indicate that HDGs encoded by viruses make a significant contribution to hydrocarbon fluxes and budgets, underscoring the necessity of incorporating virus-mediated processes into future biogeochemical cycle assessments.

## Conclusion

In summary, this study provides crucial insights into the diversity, distribution, putative host, potential contribution of VHDGs. This work significantly contributes to bridging a key knowledge gap in the understanding of global hydrocarbon degradation processes.

Future studies should aim to reveal the specific metabolic pathways and regulatory mechanisms of these genes through experimental validation and metatranscriptomic analysis, and to investigate their dynamic changes under different environmental conditions. In addition, exploring the interactions between VHDG-carrying viruses and their surrounding microbial communities will provide a more comprehensive understanding of their in situ roles, which will ultimately provide key information for bioremediation strategies and the global carbon cycle.

## Material and methods

### VHDGs acquisition and validation

The Integrated Microbial Genomes and Virome (IMG/VR) database (v4) was queried for hydrocarbon degradation genes using HMM models from the CANT-HYD[24, 25]. Enzymes involved in the activation of hydrocarbon substrates in aerobic and anaerobic hydrocarbon degradation pathways of aliphatic and aromatic compounds were identified through literature search[15]. Among them, AlkB, AlmA, and Lada/B are involved in the hydroxylation of C5-C13, C20-C32, and C15-C36 alkanes under aerobic conditions, respectively. NdoB and DszC are involved in the hydroxylation and monooxygenation of aromatic hydrocarbons under aerobic conditions, respectively. EbdA and BssA are involved in the hydroxylation and monooxygenation of aromatic hydrocarbons under anaerobic conditions, respectively. To ensure that these HDGs are encoded by viruses, the results were searched using the HMM models of viral hallmark genes from the VOG database, and illegal sequences were manually removed[59]. Finally, according to the IMG document, manually select the high-confidence viruses. A total of 768 viruses encoding hydrocarbon degradation genes were obtained for further analysis.

### World map distribution of VHDGs

Virus quality information was obtained according to the IMG documentation; the ecological niche and geographic coordinates of metagenomic samples were identified using the IMG/VR classification object IDs corresponding to each VHDGs. World map was drawn using ggplot2[60].

### Taxonomic assignments and VHDG protein grouping

The ORFs of VHDGs were mapped against the references from the International Committee on Taxonomy of Viruses (ICTV) using VITAP to determine the taxonomic affiliation[34]. To construct the protein network diagram, vConTACT2 (v.0.11.3) was used to cluster VHDGs with reference sequences from ICTV[61]. Initially, Diamond Blastp (v. 0.9.14) was employed for all-to-all comparisons among all viruses, with thresholds set at an e-value below 1e-10, sequence identity over 50%, sequence coverage over 50%[62]. The obtained results were used MCL algorithm (v. 14.137)[63]. Results exhibiting similarity to VHDGs were filtered, the network was visualized using Cytoscape(v3.10.3) and colored according to family affiliation[64].

### Molecular docking simulation

The hydrocarbons involved in the docking process consist of aliphatic and aromatic hydrocarbons, including Octane, Pentadecane, Hexadecan, Dibenzothiophene, Naphthalene, Ethylbenzene and Toluene. The three-dimensional structures of petroleum hydrocarbons were downloaded from the PubChem website (https://pubchem.ncbi.nlm.nih.gov/). Prediction of 3D structures of HDGs encoded by viruses was carried out using SWISS-MODEL automated modeling server[65]. After removing irrelevant small molecules, ligands, and water molecules from the protein structure, the file is saved in PDB format using PyMOL[66]. Subsequently, AutoDock 4.2 is used to further optimize the protein structure, including energy minimization, hydrogenation, and refinement of the protein’s three-dimensional structure[67]. The optimized structure is then saved as the receptor structure file for blind docking. The Grid Parameter File (GPF) is generated by AutoDock’s autogrid module. Finally, using the numerical values from the active pocket center region, AutoDock’s genetic algorithm is employed to estimate the binding energy scores of different conformations. The binding energy of different conformations represents the energy required to form the protein-ligand complex and can be used to evaluate the binding affinity between the ligand and the protein structure. A lower binding energy indicates a better binding conformation, with the minimum binding energy representing the optimal binding conformation[26]. The best binding conformation of each HDG with hydrocarbons is visualized using PyMOL[66].

### Viral-host prediction

Host prediction was performed using the integrated virus-host predictor iPHoP, which combines multiple virus-host prediction methods, including BLAST reference host genomes, the CRISPR spacer database, the k-mer synthesis algorithm implemented in tools such as WIsH, PHP, and VHM, and protein content-based predictions implemented in RaFAH [35, 68–71]. If multiple hosts with significant scores (>= 90) are identified, the host with CRISPR matches to the virus is considered a true positive. If no CRISPR spacer matches are identified, the host with the highest iPHoP score is considered the most likely host.

### Identification of (putatively active) proviruses and lysogens

To detect proviruses in the VHDGs, we established two criteria: (1) viral contigs originating from contigs with non-viral (host) flanking sequences, or (2) viral contigs harboring lysogenic marker proteins, which include integrase, invertase, serine recombinase, and CI/Cro repressor, as previously proposed [72]. Finally, a total of 349 proviruses were identified by this screening. Accordingly, the microorganism that harbored proviruses were regarded as lysogens[47]. Active proviruses were defined by a higher abundance of provirus sequences than the microbial host (VHR > 1), indicating active DNA replication of the provirus. Likewise, the microorganism that harbored active proviruses were regarded as active lysogens[73]. The whole genome sequence of GCF_000008865.2 was obtained from NCBI(https://www.ncbi.nlm.nih.gov/). The annotation of IMGVR_UViG_2563366825_000011 was performed using interproscan(https://www.ebi.ac.uk/interpro/). The function of each ORFs was manually checked and classified according to the e-value, and the viral genemap was drawn using ggplot2[60].

### Construction of phylogenetic tree

We downloaded reference sequences of bacteria and archaea from NCBI ((https://ftp.ncbi.nlm.nih.gov/genomes/refseq/) and used the HMM model to detect every HDGs in all sequences. We used MMseqs to create a reference non-redundant database of HDGs at the amino acid level with sequence similarity <90%[74]. These sequences, along with the HDGs encoded by viruses obtained in this study, were aligned using muscle with default parameters and trimmed at >70% gaps with trimAl[75]. We constructed a maximum likelihood phylogenetic tree using IQ-TREE (v2.0.3) with the ultrafast bootstrap approximation and the suggested protein module LG+F+R10 was employed for accurate tree[76]. ITOL (Interactive Tree Of Life) was used to visualize the phylogenetic tree[77].

### 3D Structure Comparisons

Foldseek was used to align multiple predicted protein structures via the easy-search program, with alignment fidelity (fident) calculated accordingly[78]. Clustering of 3D structures for HDG was performed using anvi-matrix-to-newick in anvi’o with manual mode[79].

### Metagenomic and metatranscriptomic mapping and gene coverage ratio calculation

Based on the results of host prediction, we downloaded protein sequences of all hosts from NCBI and searched for HDGs using HMM. We manually selected the HDGs that were the same as the virus that infected it and calculated their abundance in metagenomic and metatranscriptomic database. If the host did not encode the same HDG as the virus that infected it, the abundance was recorded as 0. We utilized metagenomic data from the Tara Oceans project, including surface, mesopelagic, deep chlorophyll maximum layer, and Mariana Trench samples[47, 80]. Additionally, metatranscriptomic data were derived from corresponding Tara Oceans datasets, as well as freshwater, microbial community, and sediment samples affected by methane pollution, obtained from the NCBI[80]. The virus:total gene coverage ratio was calculated by the HDG encoded by virus coverage values divide the virus and their host summed HDG coverage values. The gene coverage for each HDG was calculated by “jgi_- summarize_bam_contig_depths” command within metaWRAP (v1.0.2). The HDGs expression level in Reads Per Kilobase per Million mapped reads (RPKM) was calculated by normalizing the sequence depth (per million reads) and the length of the gene (in kilobases).

## Acknowledgements

We thank for the support of the high-performance servers of Center for High Performance Computing and System Simulation, Laoshan Laboratory (Qingdao), the Marine Big Data Center of Institute for Advanced Ocean Study of Ocean University of China, the High-Performance Biological Supercomputing Center at the Ocean University of China, and the IEMB-1, a high-performance computing cluster operated by the Institute of Evolution and Marine Biodiversity.

## Funding

This work was supported by the Laoshan Laboratory (No. LSKJ202203201), the Natural Science Foundation of China (No. 42120104006 and 42176111), the Ocean Negative Carbon Emissions (ONCE), the Fundamental Research Funds for the Central Universities (202172002, 201812002, 202072001 and Andrew McMinn), and the China Postdoctoral Science Foundation (2025M770867).

## Consent for publication

As requirement of JGI Data Utilization Policy, all data used in this work is published or has been gained the permissions of related data-generating principal investigators.

## Data availability

The alkane degradation viral contigs generated in this study are available in Supplementary Table1.

## Author contributions

L.R. and K.Z., performed research; K.Z curated the data; L.R. wrote the original manuscript; H.W., Z.W. and Y.L. contributed analysis tools and computing scripts; X.Z. analyzed data; Y.D H.S. provided background investigation; Y.L. X.D. A.M. and M.W. designed, administrated and supervised this project, reviewed the manuscript, and provided funding.

## Notes

### Competing Interest Statement

The authors have declared no competing interest.

